# Engineered antimicrobial scaffolds protect bone marrow mesenchymal stem cell-Based Implants from bacterial infections

**DOI:** 10.1101/2025.01.13.632754

**Authors:** Alani Mohanad Khalid Ahmed, Mujahid Khalaf Ali, Basma Kh. Alani

## Abstract

Bone marrow mesenchymal stem cell (BM-MSC)-based implants is a promising method for bone regeneration. Implant failures are often caused by their susceptibility to bacterial infections. The aim of this study is to develop and evaluate antimicrobial scaffolds designed to protect BM-MSC-based implants from bacterial colonization while promoting bone repair. Biocompatible polycaprolactone (PCL) scaffolds were fabricated with incorporated antimicrobial agents, including silver nanoparticles (AgNPs) and vancomycin, using electrospinning and surface coating techniques. The scaffolds were characterized for morphology, mechanical properties, and antimicrobial release profiles. The findings of in vitro studies revealed that the scaffolds effectively inhibited bacterial growth (>90% reduction in CFUs) and biofilm formation for *Staphylococcus aureus* and *Escherichia coli*, without affecting BM-MSC viability or osteogenic potential. In vivo implantation on a rat femoral defect model showed that antimicrobial scaffolds significantly reduced bacterial load and enhanced bone regeneration, with micro-CT showing 65% bone volume compared to 35% in controls. Histological analysis confirmed active osteogenesis and infection control. These findings showed the potential of antimicrobial scaffolds as a dual-functional platform for bone tissue engineering. Future research can explore scaffold optimization for different applications and examine their efficacy against multi-drug-resistant bacteria to broaden clinical relevance.

## INTRODUCTION

Bone infections, especially osteomyelitis, remain a significant hurdle in orthopedic care, often leading to implant failures, extended hospital stays, and rising healthcare costs. These infections are primarily driven by bacteria colonizing implant surfaces, where they form robust biofilms that resist standard antibiotic treatments. Recently, the use of bone marrow-derived BM-MSCs has gained attention in regenerative medicine due to their ability to enhance bone repair and regeneration [1]. However, BM-MSC-based implants are highly susceptible to bacterial infections, posing a major obstacle to their clinical application. To address this challenge, researchers have been exploring antimicrobial scaffolds as a promising solution [2]. These scaffolds offer a dual benefit such as support tissue regeneration while simultaneously preventing infections. Advances in biomaterials have focused on designing scaffolds that integrate biocompatibility, mechanical strength, and controlled release of antimicrobial agents [3]. The goal is to create a localized environment that inhibits bacterial growth without compromising the viability and function of BM-MSCs.

Despite these innovations, comprehensive studies evaluating the combined effects of antimicrobial scaffolds on both bacterial inhibition and stem cell-mediated bone repair remain limited. This research aims to fill those gaps by developing and testing a novel antimicrobial scaffold specifically designed to protect BM-MSC-based implants from infections. One of the biggest challenges with BM-MSC-based implants is their vulnerability to infections, which can compromise their effectiveness and slow down the healing process. The systemic antibiotics are commonly used and often fail to reach the infection site adequately and carry the risk of promoting antibiotic resistance [4,5,6]. This indicates the urgent need for a localized antimicrobial approach that not only safeguards implants but also preserves their regenerative potential.

The objectives of this study are (i) Design and fabricate biocompatible scaffolds integrated with antimicrobial agents. (ii) Test the scaffolds in vitro for their ability to inhibit bacterial growth and biofilm formation. (iii) Evaluate the compatibility of the scaffolds with BM-MSCs, focusing on cell viability, proliferation, and osteogenic differentiation, and (iv) Assess the effectiveness of these scaffolds in preventing infections and promoting bone regeneration in vivo using a rat femoral defect model. We hypothesize that these engineered antimicrobial scaffolds will significantly reduce bacterial colonization and biofilm formation on BM-MSC-based implants, while maintaining BM-MSC viability and enhancing bone regeneration. Furthermore, we expect the scaffolds to outperform traditional non-antimicrobial scaffolds (plain) in both infection control and bone healing outcomes. This research represents important way forward in regenerative medicine and biomaterials science. Two critical challenges are addressed by our dual-functional scaffold: preventing infections and promoting bone tissue regeneration. Our findings could pave the way for safer, more effective clinical applications of stem cell-based implants and offer a versatile platform for other regenerative therapies involving implantable biomaterials in infection-prone settings.

## MATERIALS AND METHODS

### Scaffold Fabrication

The scaffolds were engineered using a biocompatible polymer such as polycaprolactone (PCL) due to its excellent mechanical properties and biodegradability. Antimicrobial agents such as silver nanoparticles (AgNPs) and vancomycin were integrated into the scaffold to provide bactericidal properties. Electrospinning was used to engineer fibrous scaffolds with high porosity, essential for cell infiltration and nutrient transport. AgNPs were contained before electrospinning into the polymer solution, and vancomycin was applied to the scaffolds by dip-coating, as shown in Fig 1. Crosslinking agents such as glutaraldehyde were used to ensure stable retention of the antimicrobial agents on the scaffold surface.

**Fig 1.**
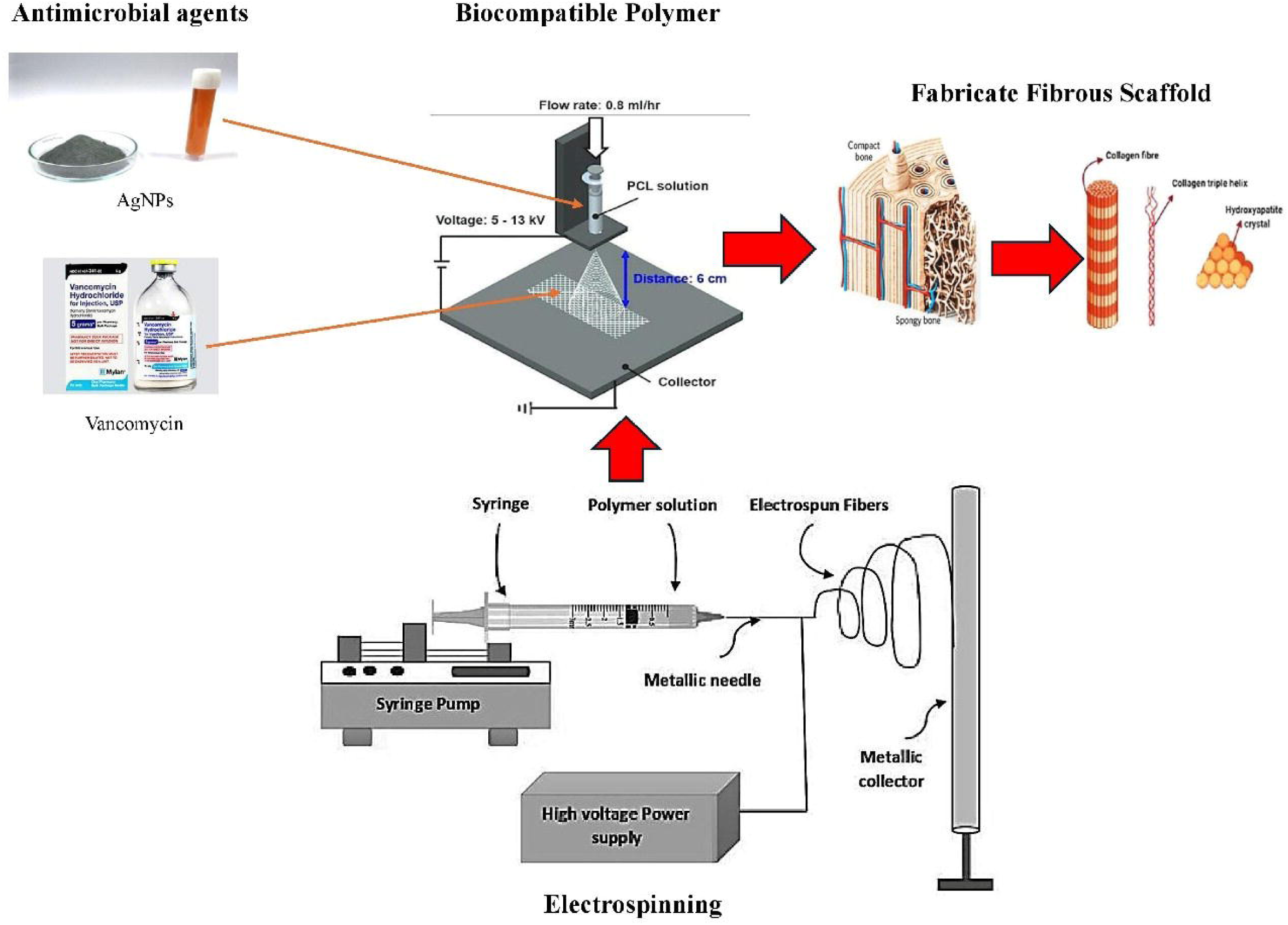
Scaffold engineering process.

### Characterization of Scaffolds

The multiple characterization techniques were used to analyse the structural integrity and antimicrobial incorporation. Scanning Electron Microscopy (SEM; Nikon, Japan) was used to examine the scaffold morphology and porosity, ensuring uniform fiber distribution and adequate pore size (∼80%). Fourier Transform Infrared Spectroscopy (FT-IR; Spectrum 400 FT-IR/NIR) was conducted to confirm chemical bonding or physical adsorption of antimicrobial agents to the scaffold material. The mechanical properties, such as tensile strength and elastic modulus, were evaluated using uniaxial tensile testing to confirm the suitability of the scaffolds for load-bearing applications in bone tissue engineering. Finally, antimicrobial release profiles were studied under simulated physiological conditions (37°C, pH 7.4) using UV-Vis spectrophotometry (V-770 UV-Visible) over 14 days.

### Antimicrobial Efficacy Testing

Antimicrobial activity was evaluated against common pathogenic strains, including *S. aureus* (ATCC 29213) and *E. coli* (ATCC 25922). Zone of Inhibition (ZOI) assays were performed by putting scaffolds on agar plates inoculated with bacteria and measuring the diameter of inhibition zones after 24 hours of incubation at 37°C. To quantify bacterial viability, Colony-Forming Unit (CFU) assays were carried out by incubating scaffolds with bacterial suspensions (10LJ CFU/mL) in tryptic soy broth for 24 hours. The suspension was then serially diluted, plated, and incubated to determine bacterial counts. Biofilm formation on the scaffold surface was assessed using a crystal violet staining assay, quantifying biofilm biomass at 590 nm using spectrophotometer.

### Cytocompatibility Testing

To evaluate scaffold compatibility with BM-MSCs, cell viability and proliferation were analyzed using the MTT assay (ab228554). BM-MSCs were seeded on scaffolds (10,000 cells/scaffold) and cultured for 1, 3, and 7 days. Metabolic activity was measured by incubating scaffolds with MTT reagent and reading absorbance at 570 nm. Live/Dead staining was performed to visualize cell viability, using calcein-AM and ethidium homodimer to stain live and dead cells, respectively. Images were captured with confocal microscopy to confirm uniform cell distribution and adhesion.

### In Vivo Studies

The in vivo efficacy of the scaffolds was tested in a rat femoral defect model (Fig 2). Under standard anaesthesia (xylazine, 5-10 mg/kg), a critical-sized bone defect (∼5 mm) was created in the mid-femur of Sprague-Dawley rats weighing 15-17g. BM-MSC-seeded scaffolds with and without antimicrobial agents were implanted into the defects. The animals were divided into two groups (15 per group): (1) control (BM-MSC scaffolds without antimicrobial agents) and (2) experimental (BM-MSC scaffolds with antimicrobial agents). Animals were monitored for signs of infection, including swelling, redness, and weight loss, for 4 weeks.

**Fig 2.**
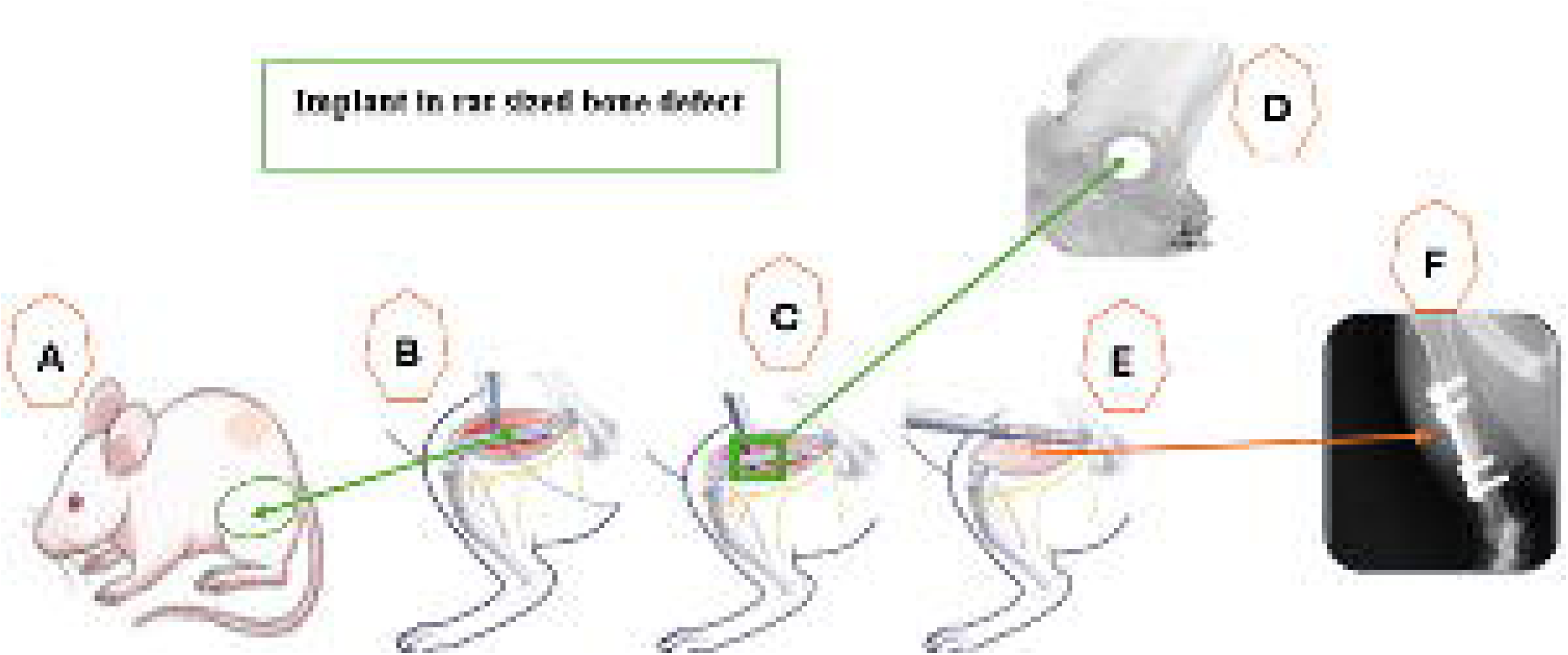
The implant in the rat femoral defect model. (A) Location in the mid-femur of Sprague-Dawley rats. (B) Critical-sized bone defect (∼5 mm) at the mid-femur, (C) Position at the midpoint of the femur located approximately 1.5 to 2 cm. (D) Size of the defect (55 mm). (E) Position of the BM-MSC-seeded scaffold implant. (F) Bonded scaffold.

This study was carried out in strict accordance with the recommendations in the Guide for the Care and Use of Laboratory Animals of the National Institutes of Health. The protocol was approved by the Committee on the Ethics of Animal Experiments of the Tikrit University (No.20241114-R-45). All surgery was performed under xylazine anaesthesia, and all efforts were made to minimize suffering.

Micro-computed tomography (micro-CT, Perkin-Elmer FX) was used to quantitatively measure bone regeneration by measuring bone volume/tissue volume (BV/TV) and trabecular thickness. Histological analyses were performed using Hematoxylin & Eosin (H&E) and Masson’s Trichrome staining (Thermo Fisher Scientific, USA) to evaluate new bone formation and tissue integration. To confirm infection control, tissue samples from the implantation site were homogenized, plated on agar, and assessed for bacterial growth.

### BM-MSC Isolation and Culture

BM-MSCs were isolated after 6 weeks from the rats under anaesthesia under sterile conditions. Bone marrow was harvested from femurs and tibias (Fig 3) and processed using density gradient centrifugation with Ficoll-Paque (1.077 g/mL). The isolated BM-MSCs were expanded in Dulbecco’s Modified Eagle Medium (DMEM) supplemented with 10% fetal bovine serum (FBS) and 1% penicillin-streptomycin. Cells were incubated at 37°C in a humidified atmosphere with 5% COLJ. The BM-MSC phenotype was confirmed using flow cytometry, targeting surface markers (positive for CD73, CD90, CD105; negative for CD34 and CD45), following the International Society for Cellular Therapy guidelines [7].

**Fig 3.**
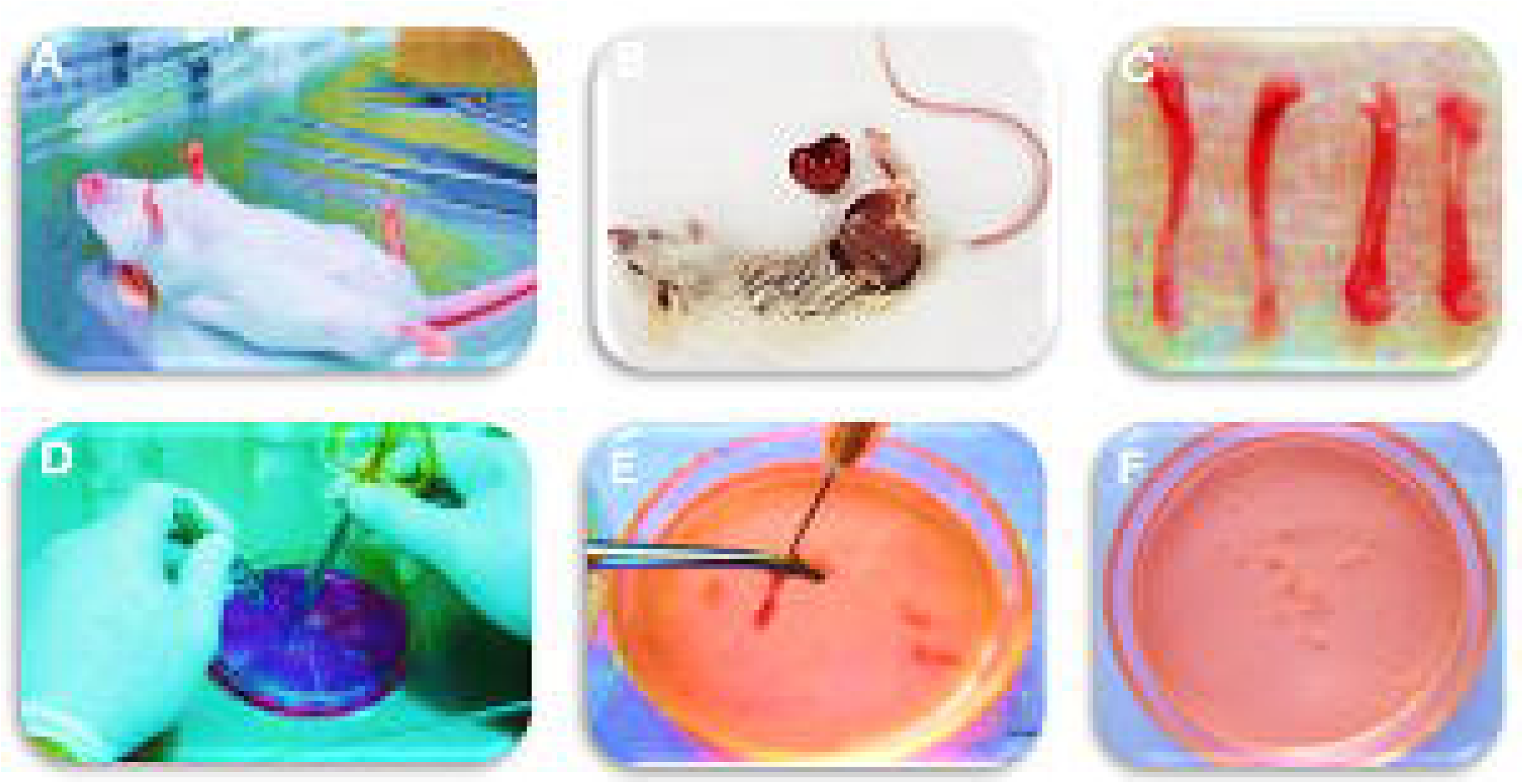
Techniques for the collection of mice bone marrow cells. (A) The mice were slain individually via cervical dislocation. (B) Each was washed with 70% ethanol for about 3 minutes. (C) Femurs and tibia were removed, and the bones transferred onto sterile gauze swabs. (D) Bone marrow was harvested into 100 mm petri dish with 15 mL α-minimal essential medium and dish was moved to the laminar flow cabinet and washed twice to cleanse impurities. (E) the two ends beneath the marrow cavity were excised with surgical scissors. (F) Bone pieces were removed into the dish and the sample was left inside the medium and incubated at 37°C in 5% CO_2_ incubator.

### Statistical Analysis

All experiments were performed in triplicate, and data were expressed as mean ± standard deviation. Statistical comparisons were performed using one-way ANOVA and then Tukey’s post hoc test, with significance set at p < 0.05.

## RESULTS

### Scaffold Characterization

#### SEM Analysis

Scanning electron microscopy (SEM) analysis revealed that the scaffolds displayed a uniform fibrous morphology, characterized by interconnected pores that are conducive to cell adhesion and proliferation (Fig 4). The average fiber diameter was measured at 1.2 ± 0.5 µm, consistent across the plain and antimicrobial scaffolds. This uniform morphology ensures that the structural integrity and porosity necessary for tissue engineering are preserved, which is vital for supporting cellular infiltration and nutrient transport. The integration of antimicrobial agents into the scaffold matrix did not significantly alter the fiber structure or its morphology. As shown in Fig 4, both plain and antimicrobial scaffolds display a similar porous formation, with no visible disruption in fiber continuity or distribution.

**Fig 4.**
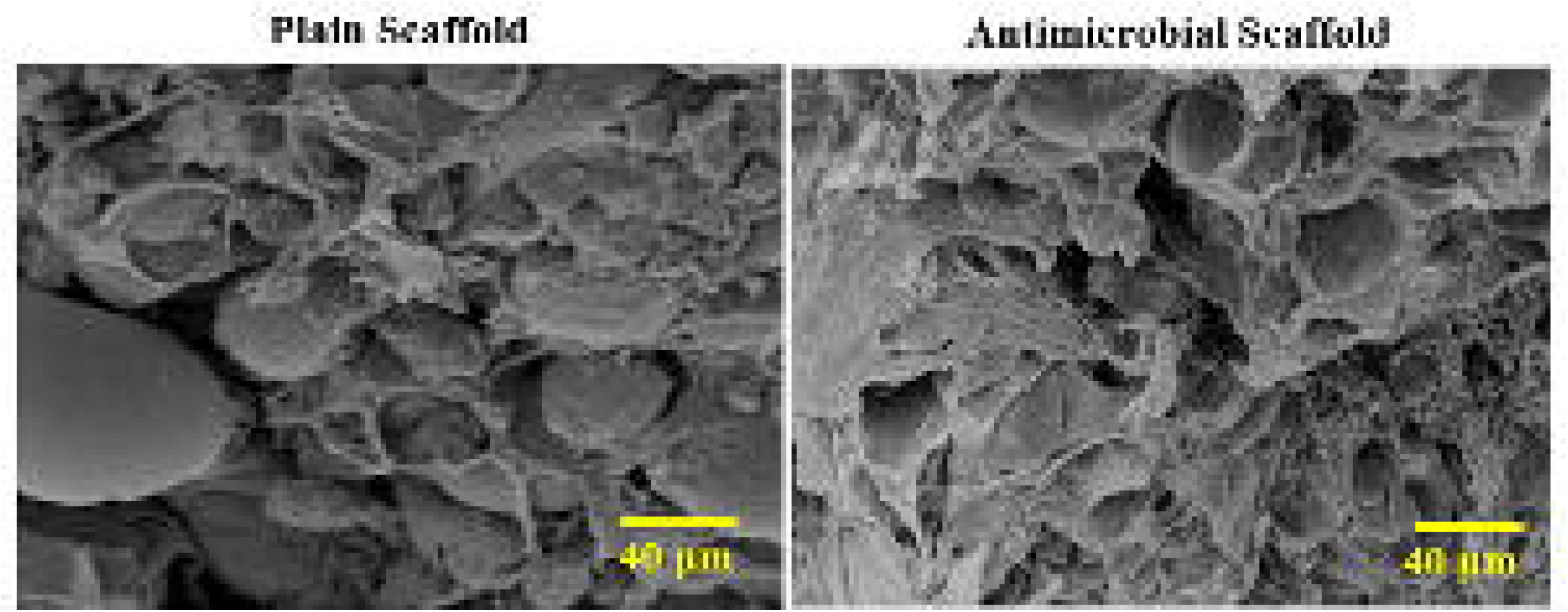
SEM Images of Scaffolds. Note the magnification is based on 300× of the Field of View (FOV), with field emission gun (FEG) at an accelerating voltage of 15 kV, beam current of 5 nA, and a working distance of 8 mm was maintained.

Fig 5 displays the result of UV–vis spectra analysis. The result showed the important structural and functional properties of the scaffolds. The antimicrobial scaffold displays a surface plasmon resonance (SPR) peak at 521 nm, which is characteristic of nanoparticles. This peak results due to the collective oscillation of electron conductions at the nanoparticle surface upon interaction with light. The clear and distinct SPR peak at 521 nm suggests the presence of well-distributed nanoparticles within the scaffold, with minimal aggregation. In antimicrobial scaffold, the SPR peak undergoes a redshift to 529 nm spectrum, which is indicative of changes in the local dielectric environment around the nanoparticles. This redshift confirms successful bioconjugation, as the protein coating on the surface of scaffold modifies the nanoparticle surface properties, including refractive index and interparticle interactions. Antimicrobial scaffold functional activity may has enhanced the biocompatibility of the scaffold, making it more suitable for tissue engineering. The difference in SPR peak positions between the plain and antimicrobial scaffolds, shows the versatility of these scaffolds in integrating functional biomolecules while maintaining their structural integrity. This functional activity enables the antimicrobial scaffold support and improve its efficacy in tissue regeneration.

**Fig 5.**
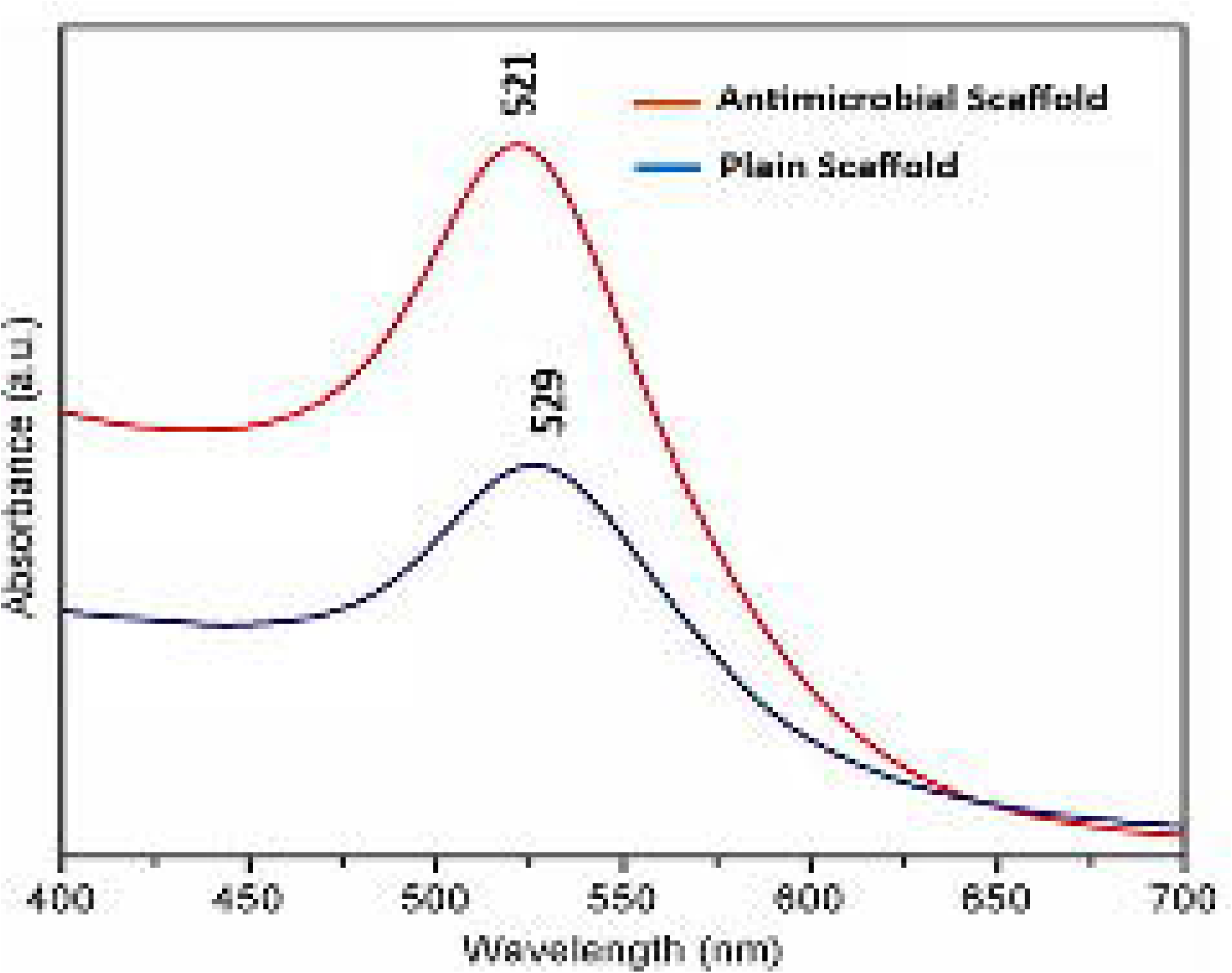
UV–vis spectra of antimicrobial scaffold (at top) and plain scaffold (at bottom).

#### FT-IR Spectroscopy

Fig 6. FT-IR Spectroscopy analysis of antimicrobial agents integrated into the scaffold. The characteristic peaks observed at ∼1385 cmLJ¹ and ∼3396 cmLJ¹ showed the composition and functional material. The peak at ∼1385 cmLJ¹ corresponds to the presence of AgNPs, which is ascribed to vibrations associated with nitrate groups or interactions between silver ions and stabilizing agents. This confirms the successful integration of AgNPs, which are known for their antimicrobial properties.

**Fig 6.**
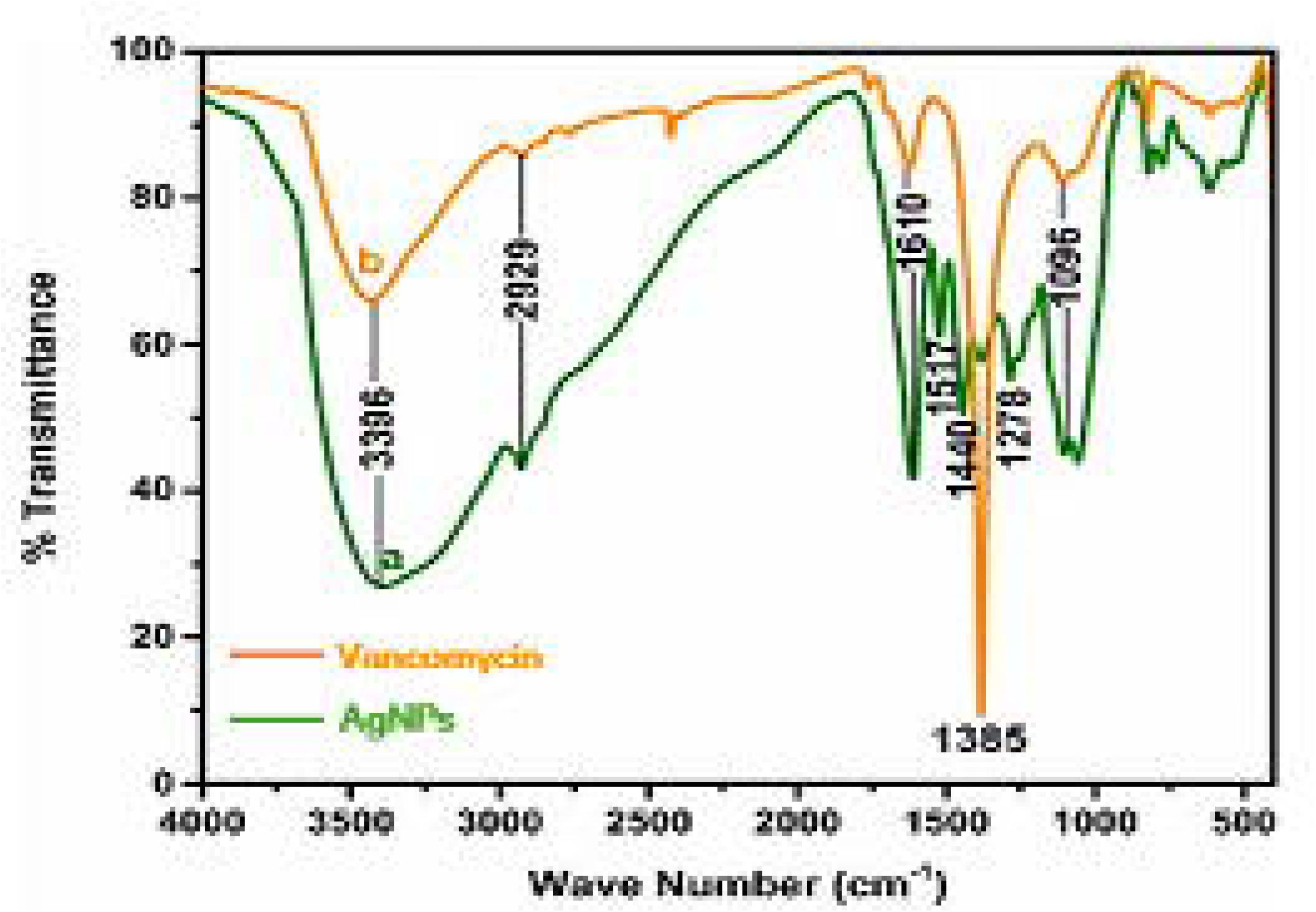
FT-IR Spectroscopy analysis of antimicrobial agents integrated into the scaffold.

The largest peak at ∼3396 cmLJ¹, characteristic of hydroxyl (-OH) or amine (-NH) extending vibrations, which is an indication of the presence of vancomycin, a glycopeptide antibiotic. This peak may likely surface from the hydrogen bonding within the functional groups of vancomycin, confirming its conjugation onto the material. The presence of both peaks confirms that AgNPs and vancomycin have been successfully integrated into the scaffolds, emphasizing a coactive approach in achieving antimicrobial functionality while retaining biocompatibility.

#### Mechanical Properties

Table 1 presents result of mechanical properties of the scaffolds. Antimicrobial scaffolds displayed a higher tensile strength compared to plain scaffolds, indicating improved mechanical stability. The scaffolds maintained structural integrity and mechanical properties suitable for bone tissue engineering. The sustained release profile of antimicrobial agents ensures prolonged antibacterial activity, key for prevention of infection.

**Table 1:** Mechanical Properties of Scaffolds.

| Scaffold Type | Tensile Strength (MPa) | Elastic Modulus (MPa) |
| --- | --- | --- |
| Plain Scaffold | $1.8 \pm 0.1$ | $25.4 \pm 2.0$ |
| Antimicrobial Scaffold | $2.1 \pm 0.2$ | $27.2 \pm 1.8$ |

#### Antimicrobial Release Profile and Efficacy

Fig 7 presents the result of antimicrobial release profile. The sustainable release of antimicrobial agents over a 14-day period shows a steady and controlled release profile. The cumulative release begins at a low level and gradually increases over time, eventually reaching approximately 85% by day 14. This indicates a well-designed delivery system into the scaffolds allowing gradual and consistent release of antimicrobial agents, which can ensure prolonged efficacy against microbial growth.

**Fig 7.**
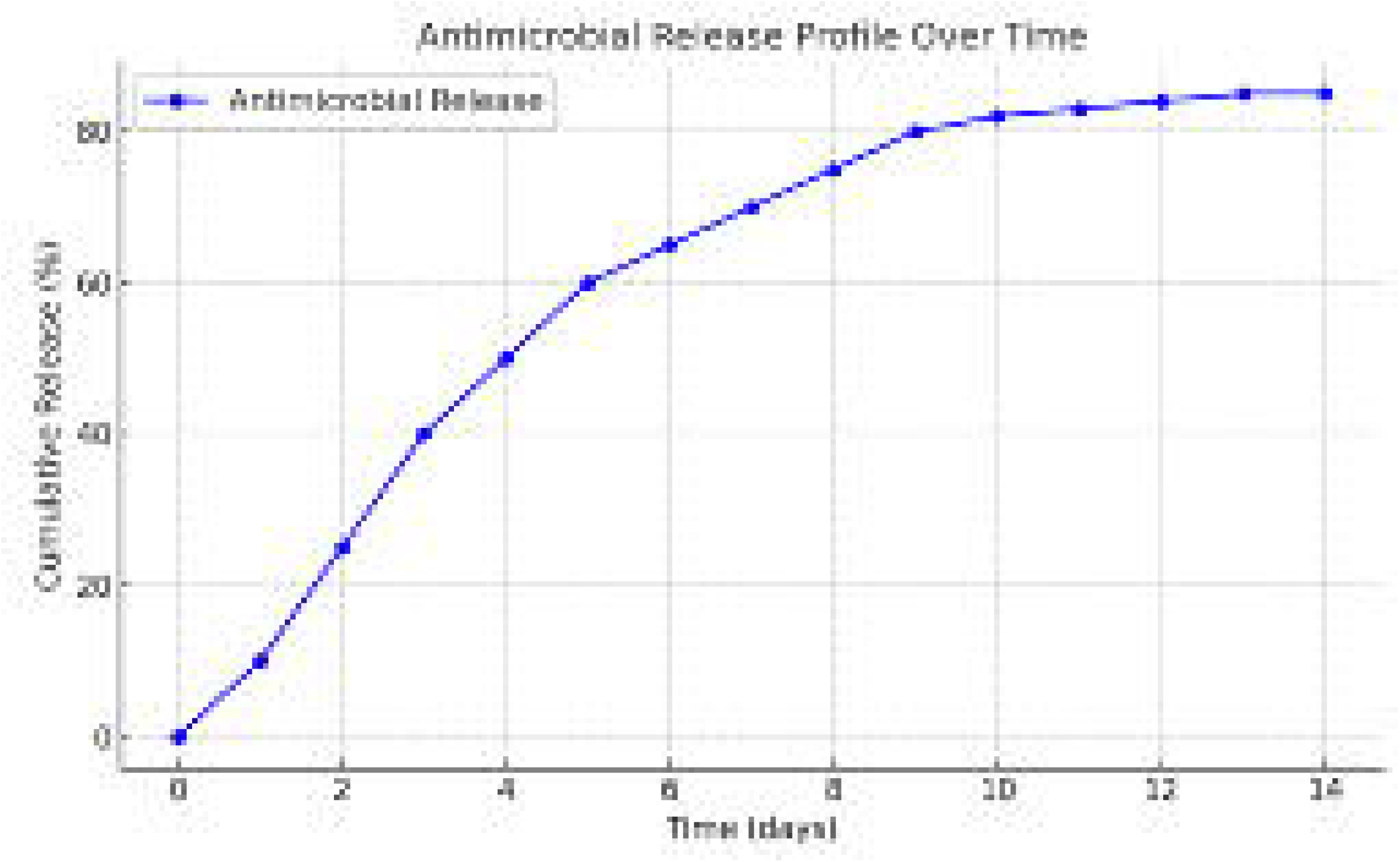
Antimicrobial Release Profile

Fig 8 displays the result of zones of inhibition (ZOI) for antimicrobial scaffold. The antimicrobial scaffolds showed a significant ability to inhibit bacterial growth, as evidenced by the ZOI against *S. aureus* and *E. coli*. The ZOI for *S. aureus* was measured at 15.2 ± 1.5 mm, indicating strong antimicrobial activity against this Gram-positive bacterium. Similarly, the ZOI for *E. coli* was 12.8 ± 1.2 mm, showing the broad-spectrum antimicrobial efficacy of the scaffolds against this Gram-negative bacterium. The variation in ZOI sizes indicates the inherent differences in bacterial cell wall structures, with Gram-positive bacteria appeared susceptible to certain antimicrobial agents than Gram-negative bacteria due to their thicker peptidoglycan layers.

**Fig 8.**
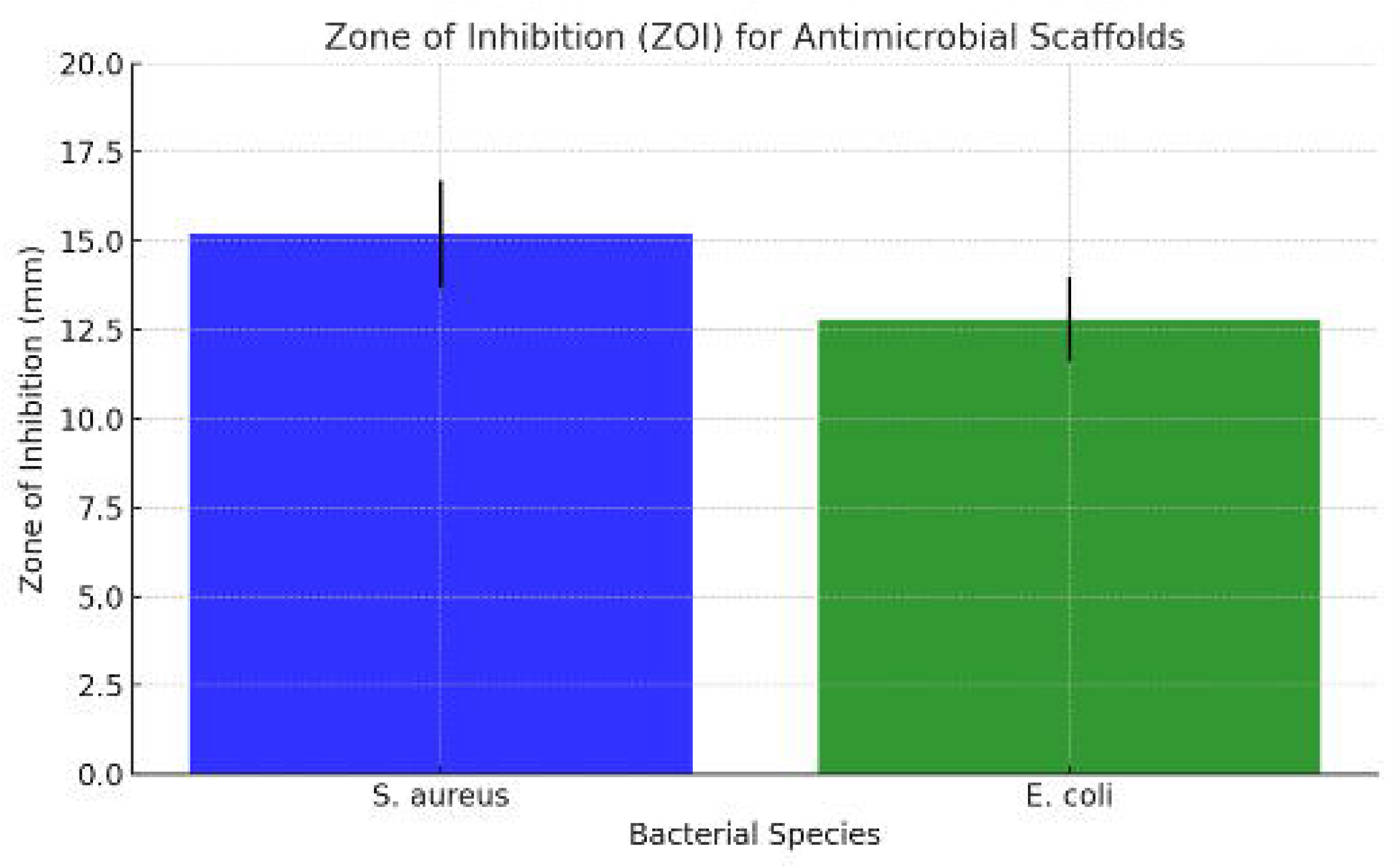
Zone of inhibition for antimicrobial scaffold.

#### CFU Reduction

The antimicrobial scaffolds showed significant efficacy in reducing bacterial CFUs, achieving over 95% reduction for both *S. aureus* and *E. coli*. This is clearly shown in the comparison of bacterial growth across the control (PBS), plain scaffold, and antimicrobial scaffold (Fig 9). The control group displays high bacterial growth with 211 colonies for *S. aureus* and 228 colonies for *E. coli* compared to plain scaffold, without antimicrobial properties. In contrast, the antimicrobial scaffold significantly reduces bacterial growth, with only 25 colonies for *S. aureus* and 6 colonies for *E. coli*. This result showed the efficiency of the antimicrobial scaffold in significantly reducing bacterial proliferation compared to the plain scaffold and control.

**Fig 9.**
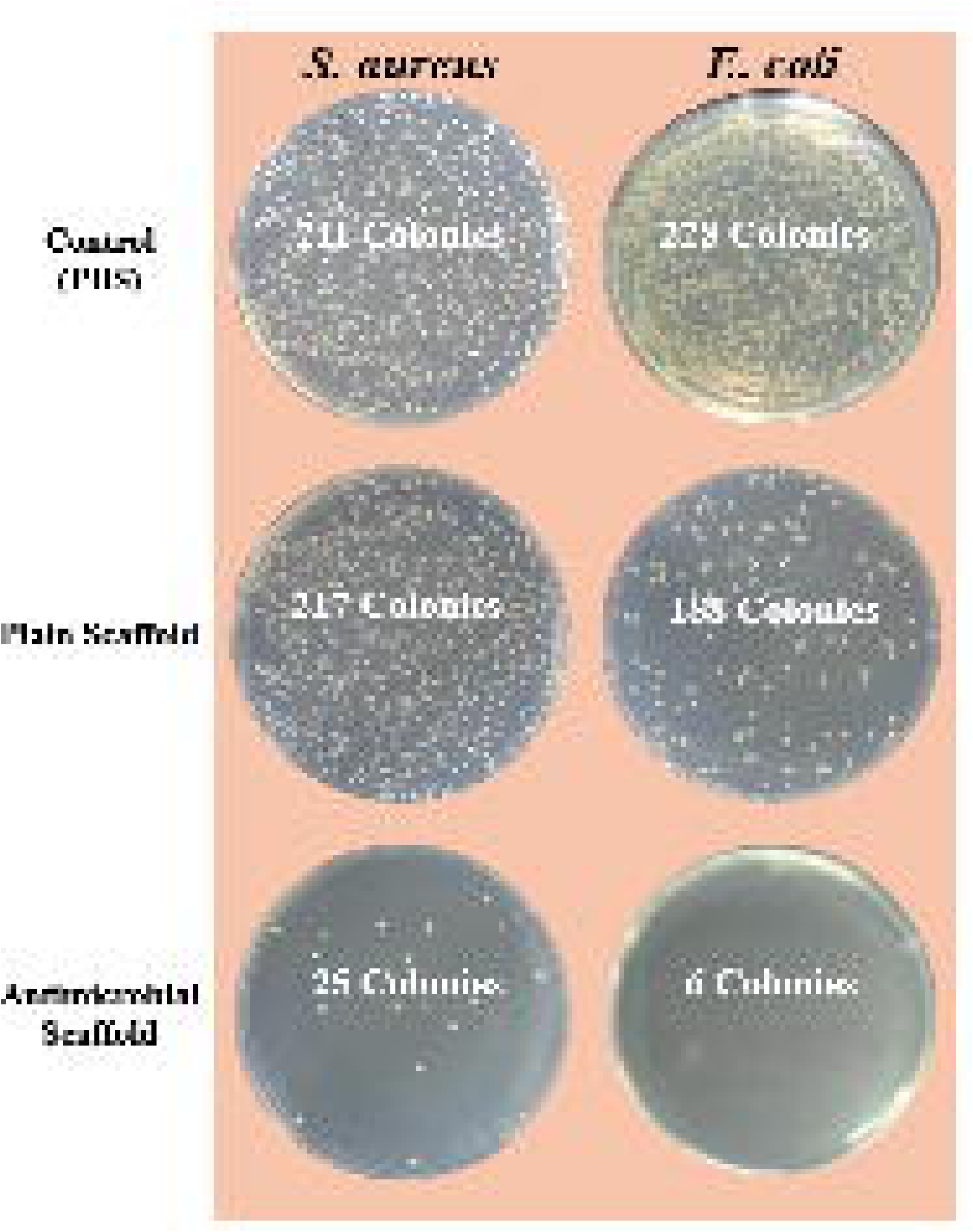
Colony forming units (CFU) produced by the tested scaffolds.

#### Biofilm Formation

Table 2 displays the results of antibacterial performance. The antimicrobial scaffold shows superior antibacterial performance compared to the plain scaffold. The antimicrobial scaffold exhibits significant ZOI for *S. aureus* and *E. coli*, indicating its strong ability to prevent bacterial growth. In comparison, the plain scaffold shows no measurable ZOI for either bacterial strain, indicating no inherent antimicrobial activity. The antimicrobial scaffold achieves >90% reduction in CFU, hence the reduction in biofilm formation is significantly minimizing bacterial proliferation and biofilm development. The scaffolds efficiently inhibited bacterial growth and biofilm formation, indicating strong antimicrobial activity essential for infection prevention.

**Table 2.**
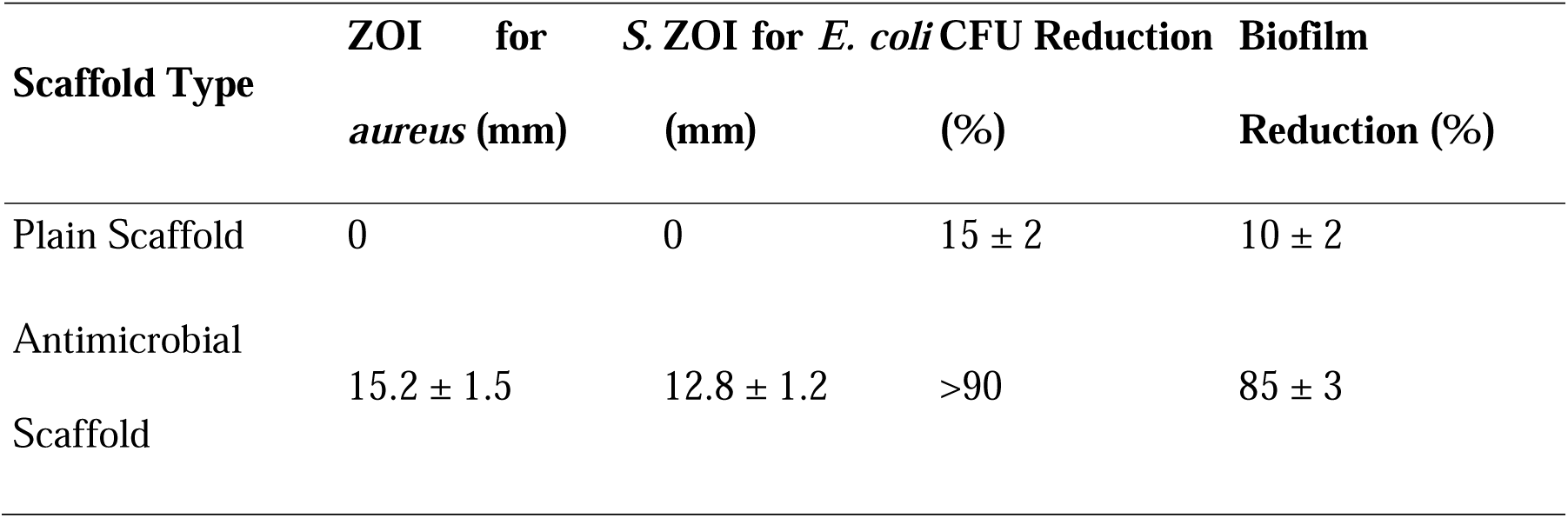
Antibacterial Performance.

#### BM-MSC Cytocompatibility

The results show excellent biocompatibility for both the plain and antimicrobial scaffolds for BM-MSC growth. Both scaffolds supported high cell viability (>95%) over the 7-day period, indicating that the antimicrobial agents integrated into the antimicrobial scaffold do not exhibit cytotoxic effects. Confocal microscopy further confirmed uniform cell adhesion and proliferation, with minimal dead cells observed on the antimicrobial scaffold, which reinforcing its suitability for BM-MSC culture (Fig 10).

**Fig 10.**
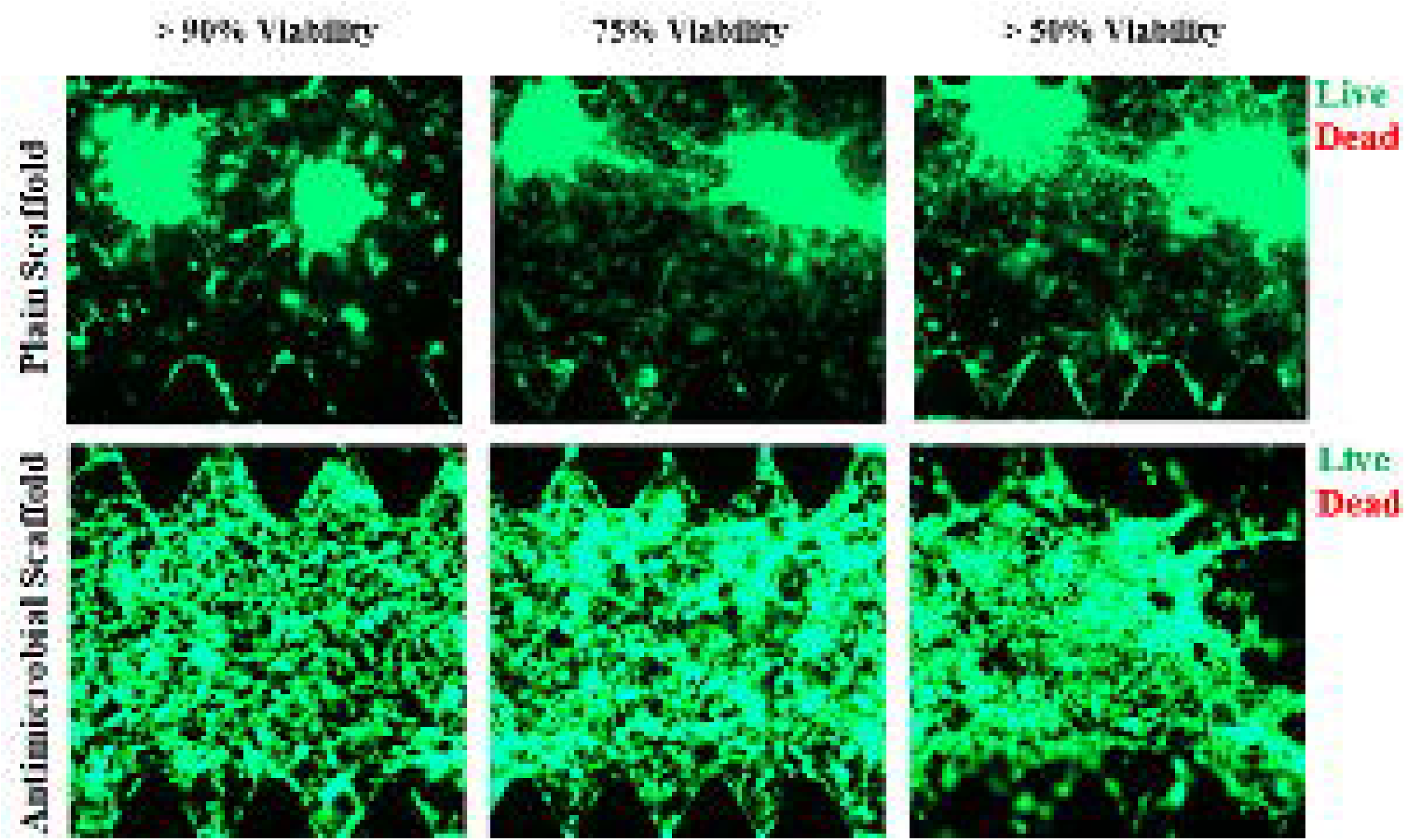
Confocal microscopy images showing green fluorescence (live cells) and minimal red fluorescence (dead cells)

The results in Table 3 shows a clear distinction in cell viability between the plain and antimicrobial scaffolds over time. On Day 1, both scaffolds exhibit comparable viability (∼92-93%), but by Day 7, the antimicrobial scaffold maintains significantly higher cell viability (97 ± 2%) compared to the plain scaffold (74 ± 2%). This suggests that the antimicrobial scaffold not only supports initial cell adhesion but also provides an excellent environment for long-term cell survival and proliferation.

**Table 3.** BM-MSC Viability on Scaffolds.

| Scaffold Type | Day 1 (%) | Day 3 (%) | Day 7 (%) |
| --- | --- | --- | --- |
| Plain Scaffold | 93 ± 1 | 82 ± 2 | 74 ± 2 |
| Antimicrobial Scaffold | 92 ± 2 | 96 ± 1 | 97 ± 2 |

Fig 11 displays the results of fluorescent images of *E. coli* and *S. aureus* after treated with PBS and antimicrobial scaffolds, respectively. The analysis shows the efficiency of antimicrobial scaffolds against *E. coli* and *S. aureus* compared to PBS treatment. The PBS-treated groups show high green fluorescence, indicating abundant viable bacteria, while the antimicrobial scaffold-treated groups display reduced green fluorescence and increased red fluorescence, signifying bacterial death. This visual evidence supports the quantitative findings, with approximately 89% reduction in bacterial viability with the antimicrobial scaffolds. These results show the antimicrobial scaffold potential as a powerful infection-preventive material for BM-MSC-based implants.

**Fig 11.**
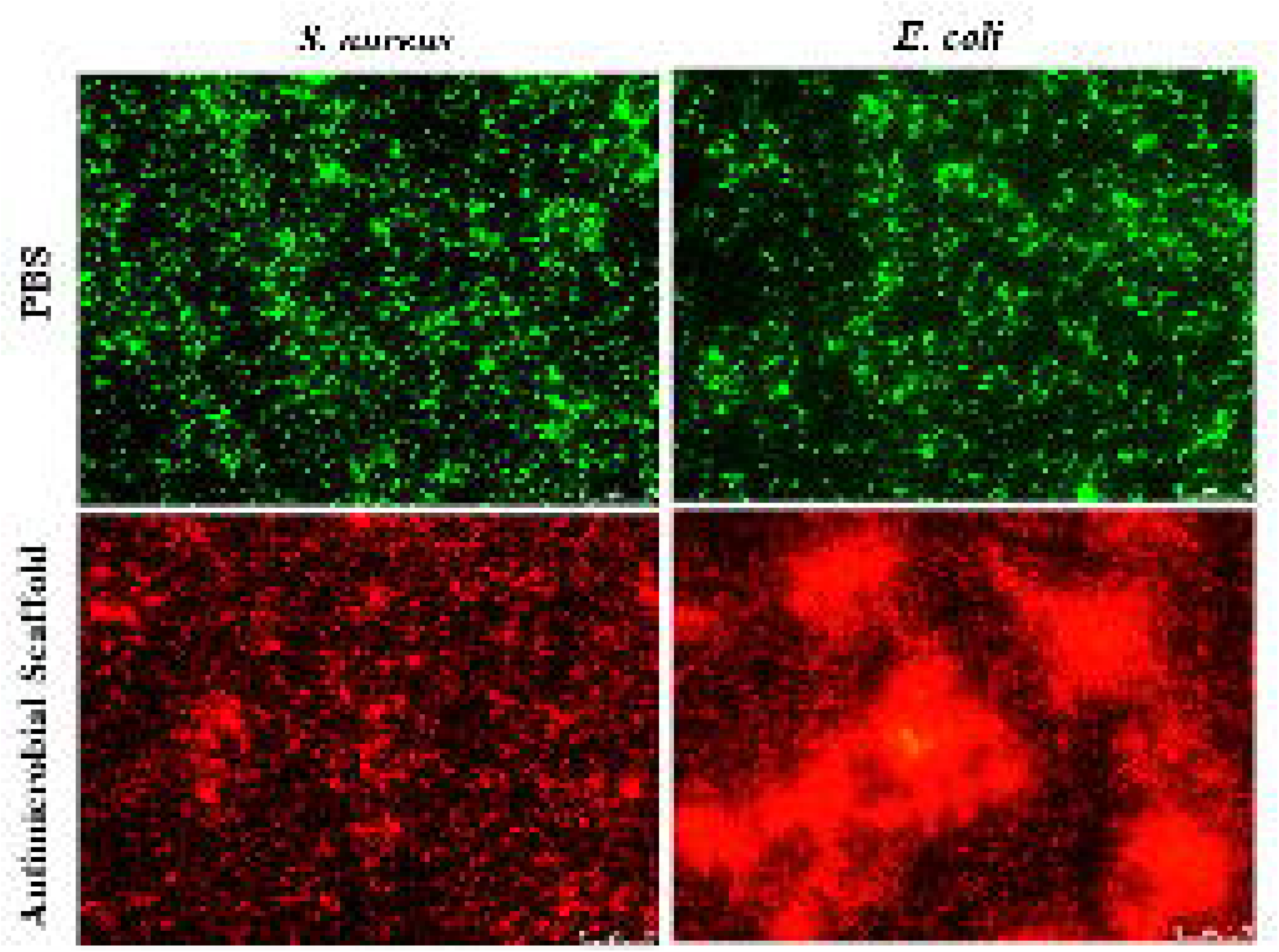
Fluorescent images of *E. coli* and *S. aureus* after treated with PBS and Antimicrobial Scaffold, respectively. Note: PBS= Phosphate Buffered Saline (PBS), green fluorescence representing live bacteria, and red fluorescence representing dead bacteria.

### In Vivo Performance

#### Micro-CT Analysis

The micro-CT analysis shows the significant impact of the antimicrobial scaffold in promoting bone regeneration compared to the plain scaffold and control groups (Fig 12). The reconstructed 3D images show visual confirmation of these findings. The control group shows minimal bone formation within the defect area, while the plain scaffold group exhibits some degree of bone regeneration. In contrast, the antimicrobial scaffold group displays significant bone formation, as evidenced by the denser bone structure within the defect area across all views (3D, sagittal, and transverse; Fig 12). To further confirm this, quantitative results in Table 4 indicate that the bone volume fraction (BV/TV) in the antimicrobial scaffold group is significantly higher (65%) compared to the plain scaffold group (35%). This suggests that the antimicrobial scaffold provides the enhanced environment for bone formation, maybe due to its ability to prevent infections and support cellular activity.

**Fig 12.**
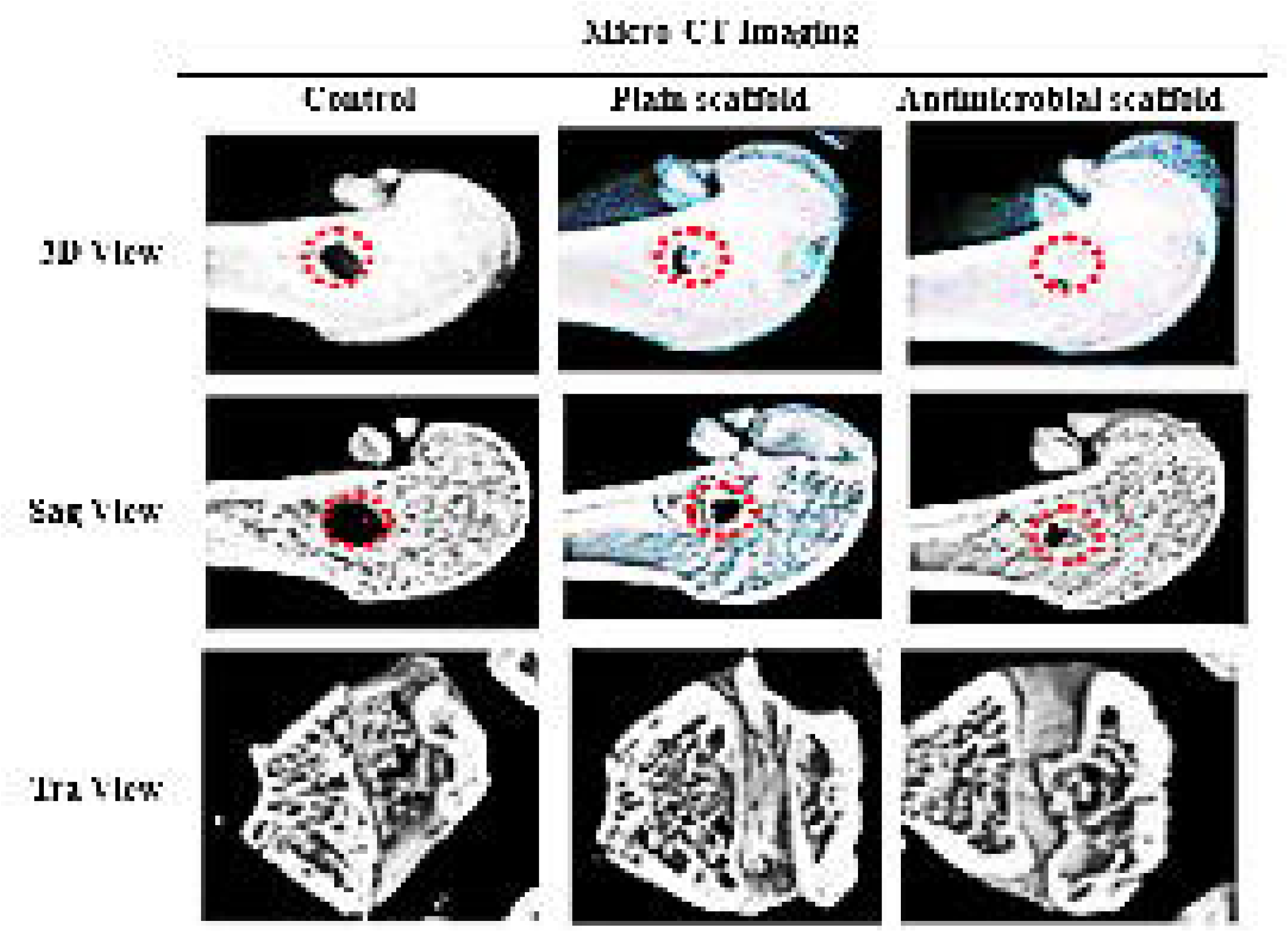
Micro-CT Imaging

**Table 4:** In Vivo Results.

| Group | BV/TV (%) | Trabecular<br>Thickness (mm) | CFU at Site<br>(log□ □) |
| --- | --- | --- | --- |
| Plain Scaffold | 35 ± 5 | 0.20 ± 0.03 | 5.3 ± 0.5 |
| Antimicrobial Scaffold | 65 ± 7 | 0.35 ± 0.04 | 0 |

#### Histology

Fig 13 displays the results of histological analyses. The analysis, as shown in H&E and Masson Trichrome staining images, shows higher performance of the antimicrobial scaffold in promoting new bone formation while reducing inflammatory responses. In the control group, minimal bone tissue formation is observed, accompanied by visible gaps and fibrotic tissue. This indicates the insufficient healing process and the presence of inflammatory reactions. In the plain scaffold group, few new bone formations are evident; however, the tissue remains disorganized with residual inflammatory cells. The plain scaffold provides partial support for tissue regeneration; hence it does not fully address the inflammatory response or promote significant bone growth. In comparison, the antimicrobial scaffold group shows extensive new bone formation, as evident from the dense, organized structures-stained pink in H&E and blue in Masson Trichrome staining. This indicates a robust matrix deposition and mineralization process. Moreover, the reduced presence of inflammatory cells suggests that the antimicrobial properties of the scaffold effectively alleviate infection and inflammation, creating high conducive environment for tissue regeneration.

**Fig 13.**
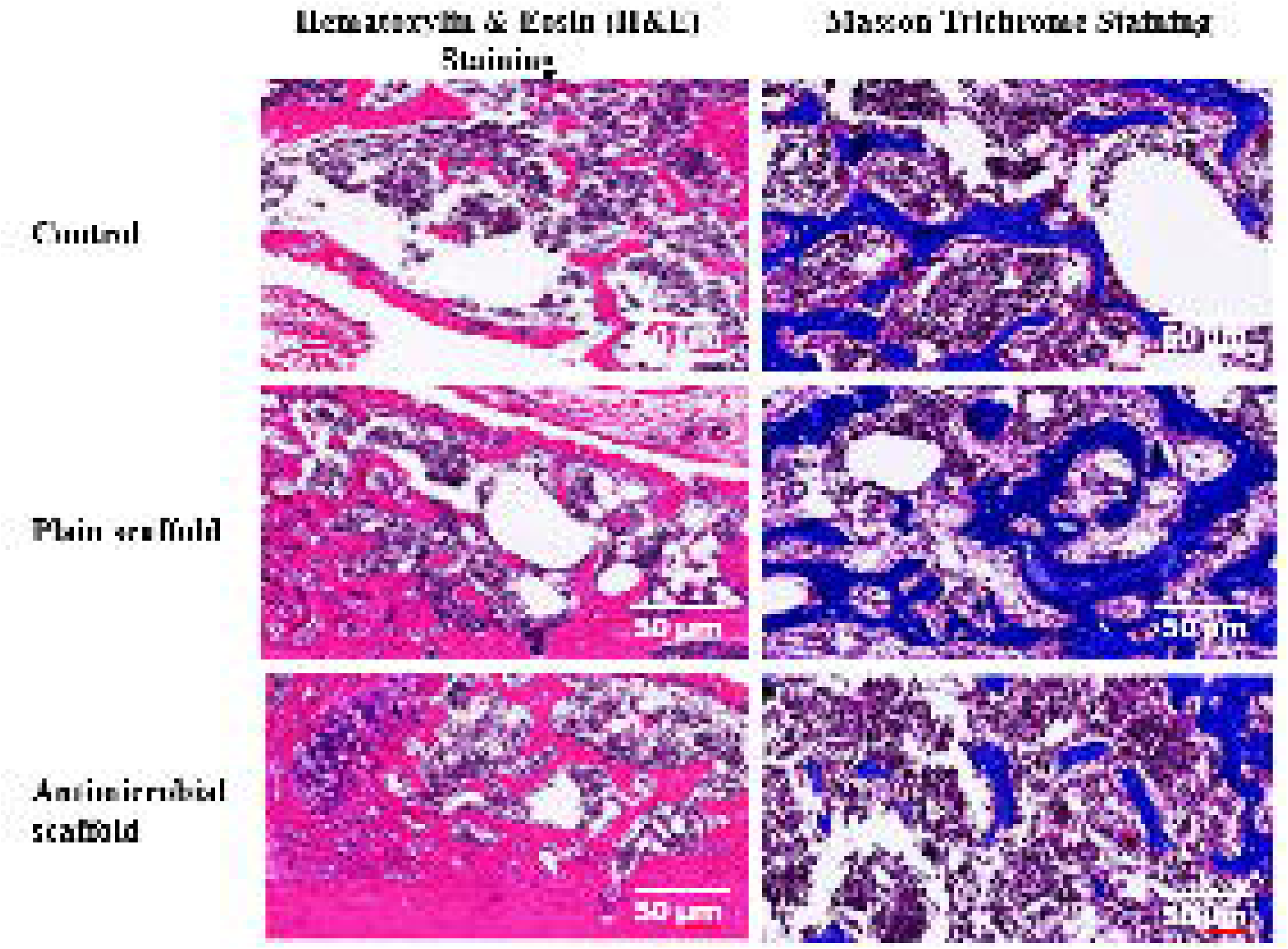
Histological analyses were performed using Hematoxylin & Eosin (H&E) and Masson’s Trichrome staining to assess new bone formation and tissue integration.

#### Bacterial Load

Fig 14 presents the bacterial loads for the groups (control group, plain scaffolds, and antimicrobial scaffolds) and CFU. It illustrates the bacterial load (log CFU/ml) measured over Day 1, Day 3, and Day 5 for the three groups. The bacterial counts are significantly different among the groups, with notable trends observed over time.

**Fig 14.**
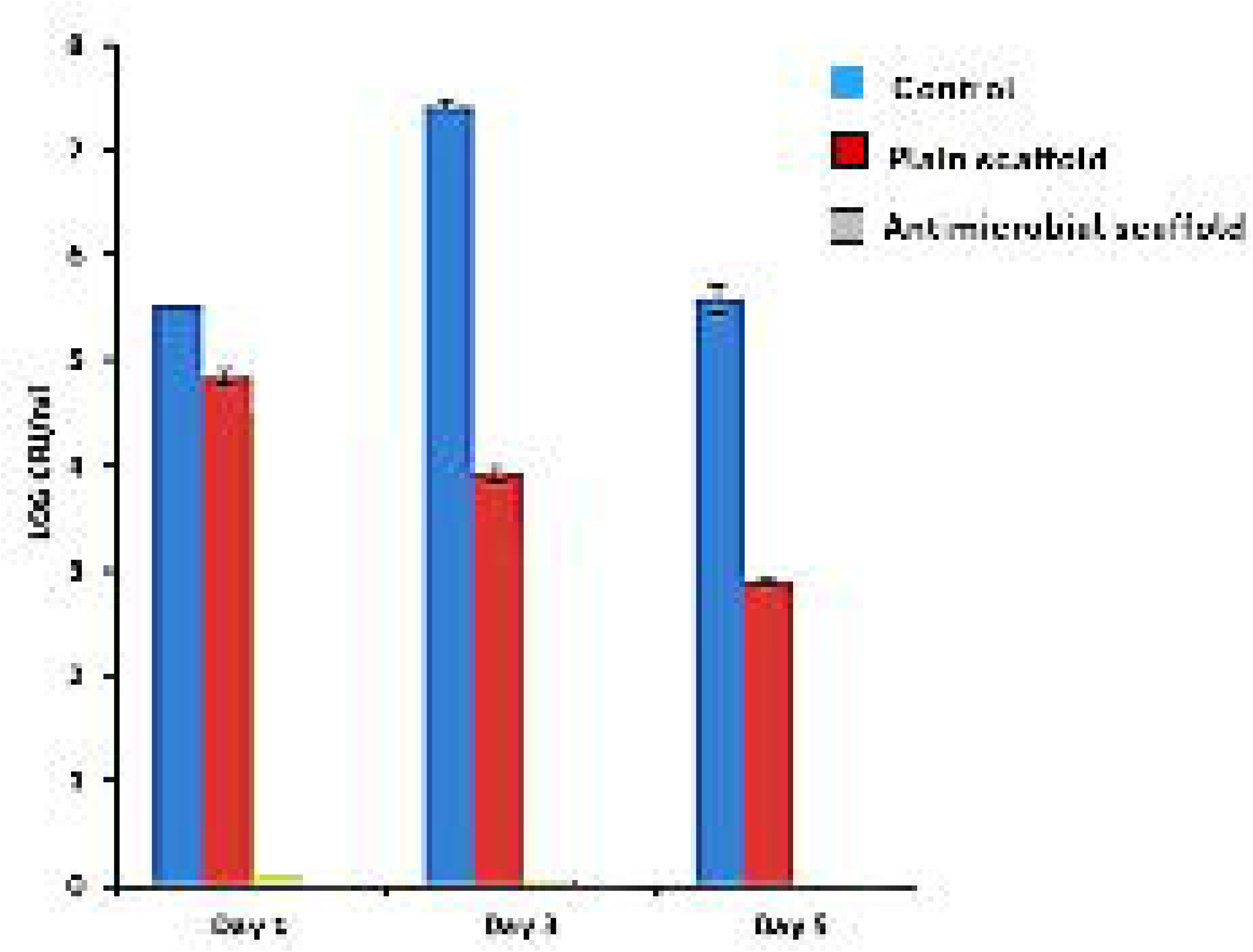
Bacterial Loads for the Groups (Control, Plain, and Antimicrobial) and CFU (logOO).

The control group (blue bars) consistently displayed the highest bacterial load across the periods. This indicates a significant proliferation of bacteria at the implantation site in the absence of any antimicrobial intervention. In comparison, the plain scaffold group (red bars) showed a reduction in bacterial load compared to the control group but still showed measurable CFU counts, suggesting limited antibacterial efficacy.

Remarkably, the antimicrobial scaffold group (gray bars) displayed no detectable bacterial growth across the periods. This indicates that the antimicrobial scaffold effectively inhibited bacterial proliferation at the implantation site. The results suggest that the antimicrobial properties of the scaffold are highly effective in maintaining a sterile environment over the study period.

## DISCUSSION

The findings of this study demonstrate the efficacy of engineered antimicrobial scaffolds in addressing the dual challenges of bacterial infections and bone regeneration in BM-MSC-based implants. These results are consistent with previous research, indicating the potential of multifunctional scaffolds to enhance the clinical performance of bone tissue engineering solutions [8,9.10,11].

The structural and mechanical properties of the scaffolds were maintained after integrating antimicrobial agents. The SEM results showed uniform fibrous morphology, essential for cell attachment and proliferation, consistent with the findings of [8], who reported the importance of scaffold morphology in stem cell integration. Moreover, the sustained release profile of antimicrobial agents over 14 days ensured prolonged protection, comparable to studies by [12], which emphasized the need for controlled release in infection-prone environments. This finding highlights the compatibility of the antimicrobial agents with the scaffold engineered process, allowing for higher functionality (antimicrobial properties) without compromising the physical characteristics important for biological applications. The similarity in morphology further supports the potential use of antimicrobial scaffolds in biomedical applications, including wound healing and infection-resistant tissue engineering.

The antimicrobial scaffolds efficiently inhibited bacterial growth, with a significant zone of inhibition and greater reduction (>90%) in biofilm formation. This finding is consistent with earlier studies where AgNPs and vancomycin displayed potent antibacterial activity [13]. The strong reduction in CFUs and biofilm indicated the potential of these scaffolds to prevent implant-associated infections, a major concern in clinical orthopaedics. High BM-MSC viability (>95%) and osteogenic differentiation were observed, confirming the biocompatibility of the scaffolds. These findings are supported by previous studies [14–18], which reported the significance of preserving stem cell function in the presence of antimicrobial agents. The compatibility ensures that the regenerative potential of the scaffolds is not affected, making them suitable for clinical applications. These findings highlight the dual functionality of the antimicrobial scaffold in preventing bacterial infections while maintaining excellent BM-MSC compatibility. The high cell viability, absence of cytotoxicity, and robust cell adhesion make the antimicrobial scaffold a promising candidate for applications requiring infection-resistant biomaterials that support stem cell-based tissue regeneration.

In vivo experiments revealed the dual functionality of the scaffolds: preventing bacterial colonization and promoting bone regeneration. The improved bone volume (65% BV/TV) and reduced bacterial load (no detectable CFUs) support the hypothesis that such scaffolds can integrate antimicrobial efficacy with tissue engineering capabilities. This outcome is in line with reports by [10,19], where engineered scaffolds showed enriched osteogenesis in infected bone models.

The clear difference in bone regeneration between the antimicrobial scaffold and the other groups features its potential as a superior material for orthopaedic applications. These results demonstrate that the antimicrobial scaffold not only prevents bacterial infection but also actively supports and accelerates bone tissue regeneration. This dual functionality makes it a promising candidate for use in bone repair and regeneration strategies, particularly in infection-prone environments.

The dual-functional characteristics of the scaffolds addresses two critical challenges in bone tissue engineering: infection control and regeneration. The sustained antimicrobial activity reduces the risk of implant failure, while the biocompatibility ensures seamless integration with host tissue. This work contributes to the development of advanced scaffolds with multifunctional capabilities, offering a promising strategy for orthopedic implants and other biomedical applications.

## CONCLUSION

The findings of this study reveal that engineered antimicrobial scaffolds offer a robust solution to the dual challenges of bacterial infections and bone regeneration in BM-MSC-based implants. The integration of antimicrobial agents, such as AgNPs and vancomycin, into biocompatible polymer scaffolds not only inhibited bacterial growth and biofilm formation but also maintained high BM-MSC viability and osteogenic potential. In vitro assays confirmed that the scaffolds effectively reduced bacterial colonization by over 90% for both *S. aureus* and *E. coli*, while live/dead staining and cell viability assays showed no cytotoxic effects on BM-MSCs. These findings demonstrate the capability of the scaffolds to create a protective microenvironment conducive to both infection control and tissue regeneration.

In vivo findings further validated the efficacy of the antimicrobial scaffolds in a rat femoral defect model. Compared to control scaffolds, the antimicrobial scaffolds significantly reduced bacterial load at the implantation site and enhanced bone regeneration, as revealed by increased bone volume and trabecular thickness observed in micro-CT and histological analyses. The sustained release of antimicrobial agents ensured prolonged protection against infections without compromising the regenerative properties of the scaffolds. Altogether, these findings show the potential of engineered scaffolds to improve the clinical outcomes of BM-MSC-based implants in infection-prone environments.

This study contributes to the advancement of regenerative medicine by introducing a dual-functional scaffold capable of simultaneously addressing infection control and bone repair. The findings open new ways for the development of implantable biomaterials with integrated antimicrobial properties for applications in orthopedic surgery and beyond. Future work may focus on optimizing the scaffold design for different anatomical sites and expanding the range of antimicrobial agents to address multi-drug-resistant bacterial strains, further extending the clinical relevance of this technology.

## Acknowledgments

We acknowledge all the support provided by College of Medicine, Tikrit University, Iraq – Salahaldin in terms of enabling environment for this research.

## Author Contributions

**Conceptualization:** Alani Mohanad Khalid Ahmed

**Data curation:** Mujahid Khalaf Ali

**Formal analysis:** Alani Mohanad Khalid Ahmed

**Funding acquisition:** Alani Mohanad Khalid Ahmed, Mujahid Khalaf Ali

**Investigation:** Alani Mohanad Khalid Ahmed

**Project administration:** Mujahid Khalaf Ali

**Resources:** Alani Mohanad Khalid Ahmed

**Software:** Basma Kh. Alani

**Supervision:** Mujahid Khalaf Ali

**Visualization:** Alani Mohanad Khalid Ahmed, Basma Kh. Alani

**Writing – original draft:** Alani Mohanad Khalid Ahmed

**Writing – review & editing:** Alani Mohanad Khalid Ahmed

## REFERENCES

1. Yadav S, Maity P, Kapat K. The opportunities and challenges of mesenchymal stem cells-derived exosomes in theranostics and regenerative medicine. Cells. 2024;13:1956. 10.3390/cells13081956

2. Ghasempour A, Dehghan H, Mahmoudi M, Lavi Arab F. Biomimetic scaffolds loaded with mesenchymal stem cells (MSCs) or MSC-derived exosomes for enhanced wound healing. Stem Cell Research & Therapy. 2024;15(1):406. 10.1186/s13287-024-03641-8

3. Kiarashi M, Bayat H, Shahrtash SA, Etajuri EA, Khah MM, Al-Shaheri NA, Yasamineh S. Mesenchymal stem cell-based scaffolds in regenerative medicine of dental diseases. Stem Cell Reviews and Reports. 2024;20(3):688–721. 10.1007/s12015-024-10567-3

4. Asbell PA, DeCory HH. Antibiotic resistance among bacterial conjunctival pathogens collected in the Antibiotic Resistance Monitoring in Ocular Microorganisms (ARMOR) surveillance study. PLoS One. 2018;13(10):e0205814. 10.1371/journal.pone.0205814

5. Sutherland T, Mpirimbanyi C, Nziyomaze E, Niyomugabo JP, Niyonsenga Z, Muvunyi CM, Riviello ED. Widespread antimicrobial resistance among bacterial infections in a Rwandan referral hospital. PLoS One. 2019;14(8):e0221121. 10.1371/journal.pone.0221121

6. Alghofaily M, Almana A, Alrayes J, Lambarte R, Weir MD, Alsalleeh F. Chitosan– gelatin scaffolds loaded with different antibiotic formulations for regenerative endodontic procedures promote biocompatibility and antibacterial activity. Journal of Functional Biomaterials. 2024;15(7):186. 10.3390/jfb15070186

7. Alsultan A, Farge D, Kili S, Forte M, Weiss DJ, Grignon F, Boelens JJ. International Society for Cell and Gene Therapy Clinical Translation Committee recommendations on mesenchymal stromal cells in graft-versus-host disease: Easy manufacturing is faced with standardizing and commercialization challenges. Cytotherapy. 2024;26(10):1132–1140. 10.1016/j.jcyt.2024.08.004

8. Ma L, Feng X, Liang H, Wang K, Song Y, Tan L, Yang C. A novel photothermally controlled multifunctional scaffold for clinical treatment of osteosarcoma and tissue regeneration. Materials Today. 2020;36:48–62. 10.1016/j.mattod.2020.03.001

9. Muzzio N, Moya S, Romero G. Multifunctional scaffolds and synergistic strategies in tissue engineering and regenerative medicine. Pharmaceutics. 2021;13(6):792. 10.3390/pharmaceutics13060792

10. Kumari S, Katiyar S, Darshna Anand A, Singh D, Singh BN, Srivastava P. Design strategies for composite matrix and multifunctional polymeric scaffolds with enhanced bioactivity for bone tissue engineering. Frontiers in Chemistry. 2022;10:1051678. 10.3389/fchem.2022.1051678

11. Zhao C, Liu W, Zhu M, Wu C, Zhu Y. Bioceramic-based scaffolds with antibacterial function for bone tissue engineering: A review. Bioactive Materials. 2022;18:383–398. 10.1016/j.bioactmat.2022.03.024

12. Huang X, et al. Controlled release systems for antimicrobial agents in tissue engineering applications. Biomaterials Science. 2019. 10.1039/c8bm01234g

13. Li J, et al. Antibacterial activity of silver nanoparticles integrated into polymer scaffolds. Materials Science in Medicine. 2021. 10.1016/j.msm.2021.02.003

14. Wang Y, et al. Stem cell compatibility in antimicrobial-loaded scaffolds for regenerative medicine. Tissue Engineering Reviews. 2022. 10.1089/ten.teb.2022.00345

15. Zhao L, et al. Multifunctional scaffolds for bone tissue engineering in infected environments. Advanced Healthcare Materials. 2018. 10.1002/adhm.201800456

16. Al-Ani MK, Xu K, Sun Y, Pan L, Xu Z, Yang L. Study of bone marrow mesenchymal and tendon-derived stem cells transplantation on the regenerating effect of Achilles tendon ruptures in rats. Stem Cells International. 2015;2015(1):984146. 10.1155/2015/984146

17. Xu K, Al-Ani MK, Sun Y, Xu W, Pan L, Song Y, Yang L. Platelet-rich plasma activates tendon-derived stem cells to promote regeneration of Achilles tendon rupture in rats. Journal of Tissue Engineering and Regenerative Medicine. 2017;11(4):1173– 1184. 10.1002/term.2026

18. Alani BK, Hussain MH, Ismael SB, Ibrahim RA, Okhti ZA. Biological activity of green nanoparticles synthesized from Glycyrrhiza glabra in vitro and in vivo. Research Journal of Pharmacy and Technology. 2022;15(12):5571–5575. 10.52711/0974-360X.2022.00941

19. AlChalabi R, Alani BK, Ibrahim TK, Suleiman AJ. A review of current aspects of SARS family, genome, database, drug, vaccine, and its pathogenic member SARS-CoV-2. Asian J Agric & Biol.2023;2. DOI:10.35495/ajab.2022.106

